# Modulation of *ACD6* dependent hyperimmunity by natural alleles of an *Arabidopsis thaliana* NLR resistance gene

**DOI:** 10.1101/300798

**Authors:** Wangsheng Zhu, Maricris Zaidem, Anna-Lena Van de Weyer, Rafal M. Gutaker, Eunyoung Chae, Sang-Tae Kim, Felix Bemm, Lei Li, Rebecca Schwab, Frederik Unger, Marcel Janis Beha, Monika Demar, Detlef Weigel

## Abstract

Plants defend themselves against pathogens by activating an array of immune responses. Unfortunately, immunity programs may also cause unintended collateral damage to the plant itself. The quantitative disease resistance gene *ACCELERATED CELL DEATH 6* (*ACD6*) serves as a nexus for the trade-off between growth and pathogen resistance in wild populations of *Arabidopsis thaliana.* An autoimmune allele, *ACD6*-Est, first identified in the natural accession Est-1, is found in over 10% of wild strains, even though it causes a clear fitness penalty under optimal growth conditions. There is, however, extensive variation in the strength of the autoimmune phenotype expressed by strains with an *ACD6*-Est allele, indicative of genetic modifiers. Quantitative genetic analysis suggests that the population genetic basis of *ACD6* modulation is complex, with different strains often carrying different large-effect modifiers. One modifier is *SUPPRESSOR OF NPR1*-*1*, *CONSTITUTIVE 1* (*SNC1*), located in a highly polymorphic cluster of nucleotide-binding domain and leucine-rich repeat (NLR) immune receptor genes, which are prototypes for qualitative disease resistance genes. Allelic variation at *SNC1* correlates with *ACD6*-Est activity in multiple accessions, and a common structural variant affecting the NL linker sequence can explain differences in SNC1 activity. Taken together, we find that an NLR gene can mask the activity of an *ACD6* autoimmune allele in natural *A. thaliana* populations, thereby linking different arms of the plant immune system.

**Author summary:** Plants defend themselves against pathogens by activating immune responses. Unfortunately, these can cause unintended collateral damage to the plant itself. Nevertheless, some wild plants have genetic variants that confer a low threshold for the activation of immunity. While these enable a plant to respond particularly quickly to pathogen attack, such variants might be potentially dangerous. We are investigating one such variant of the immune gene *ACCELERATED CELL DEATH 6* (*ACD6*) in the plant *Arabidopsis thaliana.* We discovered that there are variants at other genetic loci that can mask the effects of an overly active *ACD6* gene. One of these genes, *SUPPRESSOR OF NPR1*-*1*, *CONSTITUTIVE 1* (*SNC1*), codes for a known immune receptor. The *SNC1* variant that attenuates *ACD6* activity is rather common in *A. thaliana* populations, suggesting that new combinations of the hyperactive *ACD6* variant and this antagonistic *SNC1* variant will often arise by natural crosses. Similarly, because the two genes are unlinked, outcrossing will often lead to the hyperactive *ACD6* variants being unmasked again. We propose that allelic diversity at *SNC1* contributes to the maintenance of the hyperactive *ACD6* variant in natural *A. thaliana* populations.

## Introduction

Plants rely on a sophisticated immune system to defend themselves against pathogens. A central challenge for plants is how to achieve a fast, effective response upon pathogen attack, while at the same time preventing spontaneous firing of the signaling machinery in the absence of danger [1]. Inappropriate activation of immune signaling can reduce growth or even damage the plant’s own cells, while an inefficient immune response makes plants more likely to succumb to pathogen attack [2–6]. Although highly effective immune alleles may have background activity, they will nevertheless be favored when pathogen pressure is high. In contrast, in locations and years with low pathogen pressure, such alleles tend to be selected against [7]. If differently active alleles exist at the same locus, such temporal or spatial variation in pathogen pressure will maintain both types of alleles at ratios that reflect the prevalence of the different environments; this is one example of the phenomenon of balancing selection [8–10]. Very different alleles have been described for several disease resistance (R) genes of the nucleotide binding site leucine-rich repeat (NLR) class, although formal evidence for balancing selection is still rare [11–15]. NLR immune receptors detect the presence of so-called pathogen effector molecules, leading to effector-triggered immunity (ETI). NLRs do so in several different ways, and the biochemical and structural basis for effector detection, including direct interaction with effectors or with effector-modified host proteins, is increasingly well understood [16,17]. Variation in NLR immune receptors is linked to the fact that effectors are rarely essential for pathogen survival, and that even closely related pathogens can greatly vary in effector content [16,18].

In mutant screens, gain-of-function alleles have been identified for several NLR genes, based on their autoimmune phenotypes. The mutant proteins are active regardless of effector presence, and thus can increase resistance to a range of unrelated pathogens [19–26]. Whether highly active autoimmune alleles of NLR genes are being deployed to enhance broad-spectrum pathogen resistance in nature is unknown, but the presence of NLR alleles with modest autoimmune activity in breeding programs is well documented, and phenotypic signs of autoimmunity have been exploited to follow resistance genes in segregating crop populations (e.g., [27]).

In addition to specialized NLRs, which often engender very strong, qualitative disease resistance, plants employ genes for quantitative resistance. One example is *ACCELERATED CELL DEATH 6* (*ACD6*) [28–31]. Although *ACD6* does not encode an NLR immune receptor, the locus features extensive copy number and sequence variation in wild populations, reminiscent of what is found at many NLR loci. Importantly, natural populations segregate for *ACD6* allele classes with clear functional differences [32–34]. One class, first identified by genetic analysis of the natural Est-1 accession, protects in the greenhouse against a wide range of unrelated pathogens, from microbes to insects [32]. This unusually large benefit of the hyperactive *ACD6* allele appears to be due to enhanced and partially constitutive defense responses. On the other hand, the autoimmunity seen in plants with the *ACD6*-Est allele substantially compromises growth under optimal conditions, reducing both plant size and the tempo with which new leaves are being produced. Nevertheless, even though *ACD6*-Est imposes a substantial penalty on plant growth under optimal conditions, it is found at a frequency of over 10% in natural populations [32–34]. Whether the growth penalties are less severe under natural conditions, and how effectively *ACD6*-Est protects from pathogens in the wild, is not known, but that the frequency of functionally distinct *ACD6* alleles differs between local populations is suggestive of a role of *ACD6* in local adaptation [33,34].

*ACD6* encodes a transmembrane protein with intracellular ankyrin repeats, which positively regulates cell death and defense, acting in part via the immune hormone salicylic acid (SA) and the SA transducer *NONEXPRESSER OF PR GENES 1* (*NPR1*) [28,29]. The precise mechanism of ACD6 biochemical action remains enigmatic, but ACD6 is found in a complex with several immune-related proteins, including pathogen-associated-molecular-pattern (PAMP) receptors, indicating a role for ACD6 in PAMP triggered immunity (PTI) [30,31]. While PTI and ETI have often been considered as different arms of the plant immune system, the distinction between ETI and PTI is becoming more and more blurred, not only because of substantial overlap in downstream responses, but also because not all effectors are highly variable, and not all PAMPs are highly conserved [35].

Here, we report that natural alleles at several independent loci can modulate the activity of the *ACD6*-Est autoimmune allele. We have characterized one locus, encoding the NLR protein SUPPRESSOR OF NPR1-1, CONSTITUTIVE 1 (SNC1) in detail and show that a common structural variant affecting the NL linker explains differences in *ACD6*-Est dependent SNC1 activity. We propose that allelic diversity at the *SNC1* locus contributes to the maintenance of the *ACD6*-Est autoimmune allele in natural *A. thaliana* populations.

## Results

### Incidence of the *ACD6*-Est autoimmune allele in natural *A. thaliana* populations

Sanger sequencing analysis of a limited collection of 96 natural accessions had shown that 19, or 20%, had the *ACD6*-Est autoimmune allele [32]. To extend our knowledge of the global distribution of the *ACD6*-Est allele, we attempted to use Illumina short read data of 1,135 accessions [36,37], with the goal of ascertaining the presence of two codons that are causal for *ACD6*-Est autoimmune activity [32]. Likely because of linked sequence diversity and copy number variation, the first codon, encoding amino acid 566 in the reference sequence, GCA (Ala) in Col-0 and AAC (Asn) in Est-1, could not be confidently typed with short reads. At the second codon, encoding amino acid 634 in the reference, 823 accessions could be assigned to have either Col-0- or Est-1-like codons. Of these, 721, or 88%, had CTT (Leu), diagnostic for the Col-0 reference, and 102, or 12%, had TTT (Phe), diagnostic for Est-1, confirming that *ACD6*-Est alleles are not uncommon in the global *A. thaliana* population.

Of the 19 accessions with an *ACD6*-Est allele examined by Todesco and colleagues [32], two did not show any necrotic lesions, an obvious sign of autoimmunity. The remaining 17 accessions differed in their phenotypic severity as well, with three classified as expressing mild, five intermediate, and nine strong necrosis. We extended this analysis to 67 further accessions for which we had confirmed the presence of an *ACD6*-Est allele by Sanger sequencing **(S1 Table)**. Fourteen had no or only very mild lesions, 25 had intermediate lesions, and 27 were similarly affected as the Est-1 strain, in which this allele had been originally identified **(Fig 1A)**.

**Figure 1.**
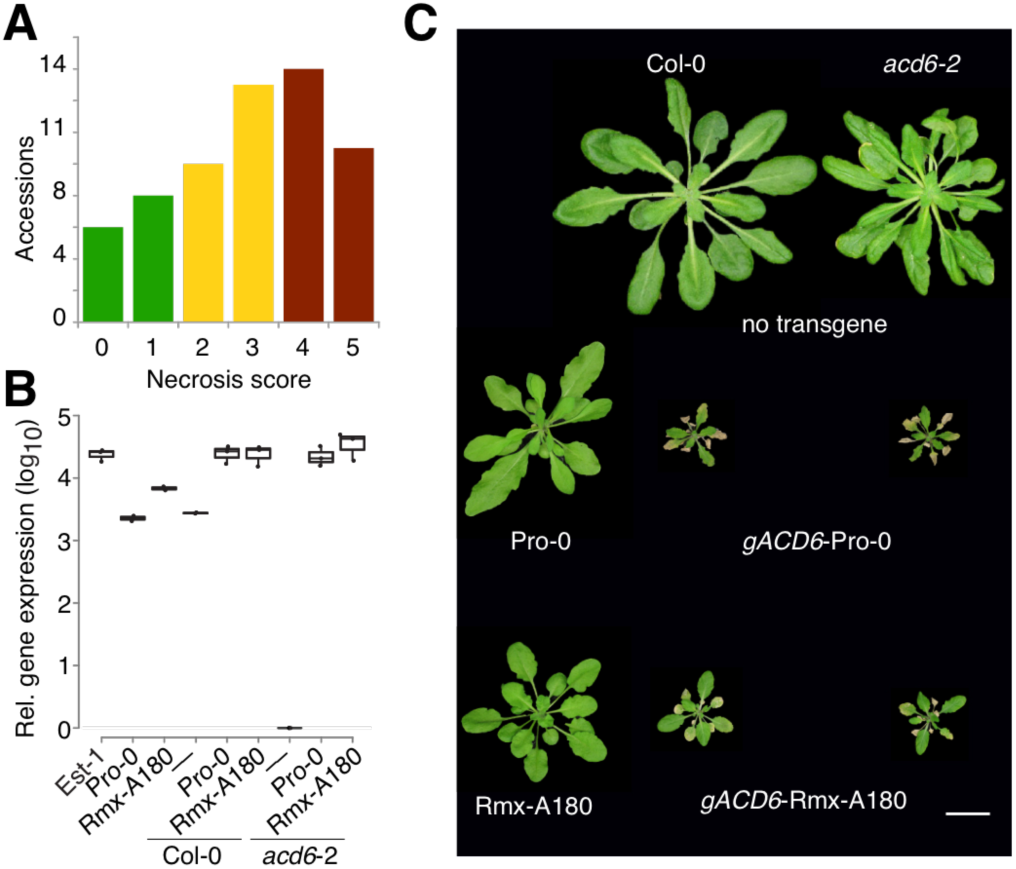
Genetic background dependence of *ACD6*-Est effect. (A) Variation of necrosis in natural accessions with *ACD6*-Est alleles. (B) Accumulation of *ACD6* mRNA in various genetic backgrounds, as measured by qRT-PCR from three biological replicates each. (C) Representative non-transgenic and *ACD6* transgenic lines at 5 weeks of age in Col-0 and *acd6*-*2* backgrounds. Pro-0 and Rmx-A180 are shown for comparison on the left. Scale bar, 10 mm.

### Evidence for extragenic suppression of *ACD6*-Est alleles

While we could confirm that there is substantial variation in the extent of autoimmune phenotypes exhibited by *ACD6*-Est carriers, this observation did not inform about the genetic cause of this variation. To determine whether this was due to to sequence differences at the *ACD6* locus itself, we functionally analyzed the *ACD6* locus in the two *ACD6*-Est accessions, originally identified as lacking lesions, Pro-0 and Rmx-A180 [32]. *ACD6* was expressed well in both accessions, indicating that suppression of the *ACD6*-Est phenotype was not due to reduced RNA accumulation **(Fig 1B)**. We introduced full-length genomic fragments from both accessions into the Col-0 reference strain, which carries a standard *ACD6* allele, and into *acd6*-*2*, a Col-0 derivative with a T-DNA insertion in *ACD6*. All four classes of transformants from 10 T_1_ lines had small rosettes and necrotic lesions, similar to what has been reported for Est-1 [32] **(Fig 1C).** Together, these results demonstrate that the *ACD6* alles from Pro-0 and Rmx-A180 have similar activity as the original *ACD6*-Est allele outside their native genetic backgrounds, suggesting that accessions such as Pro-0 and Rmx-A-180 carry extragenic suppressors of *ACD6*-Est activity.

*ACD6* acts in a feed-forward loop that regulates the accumulation of SA, a key hormone in both disease signaling and autoimmunity [38]. We therefore asked whether the suppressed autoimmunity of the *ACD6*-Est allele in Pro-0 was accompanied by reduced SA levels. Indeed, Pro-0 had much less SA than the *ACD6*-Est reference strain Est-1 or *acd6*-*1*, which carries an EMS-induced hyperactive *ACD6* allele in the Col-0 background [29]. SA levels in Pro-0 were even lower than in Col-0, which carries the standard *ACD6* allele **(Fig 2A)**. Similarly, the expression of SA responsive marker gene *PR1* was reduced in Pro-0 compared to Est-1 **(Fig 2B)**. As expected, knocking down *ACD6* with an artificial miRNA [32,39] resulted in greatly reduced SA levels in both *acd6*-*1* and Est-1, while the effect in Pro-0 was much smaller **(Fig 2A**). In agreement, knocking-down *ACD6* in Pro-0 did not substantially affect *PR1* expression either **(Fig 2B)**. Finally, to assess not only autoimmunity but also true immune responses, we measured reactive-oxygen species (ROS) accumulation after stimulation with the PAMP flg22 [40]. Est-1 responded strongly to flg22, with the ROS burst being greatly reduced by knocking down *ACD6.* Pro-0 responded much more weakly, and knocking down *ACD6* was of little consequence **(Fig 2C)**. In conclusion, these results confirmed attenuation of the effects of the hyperactive *ACD6*-Est allele in Pro-0.

**Figure 2.**
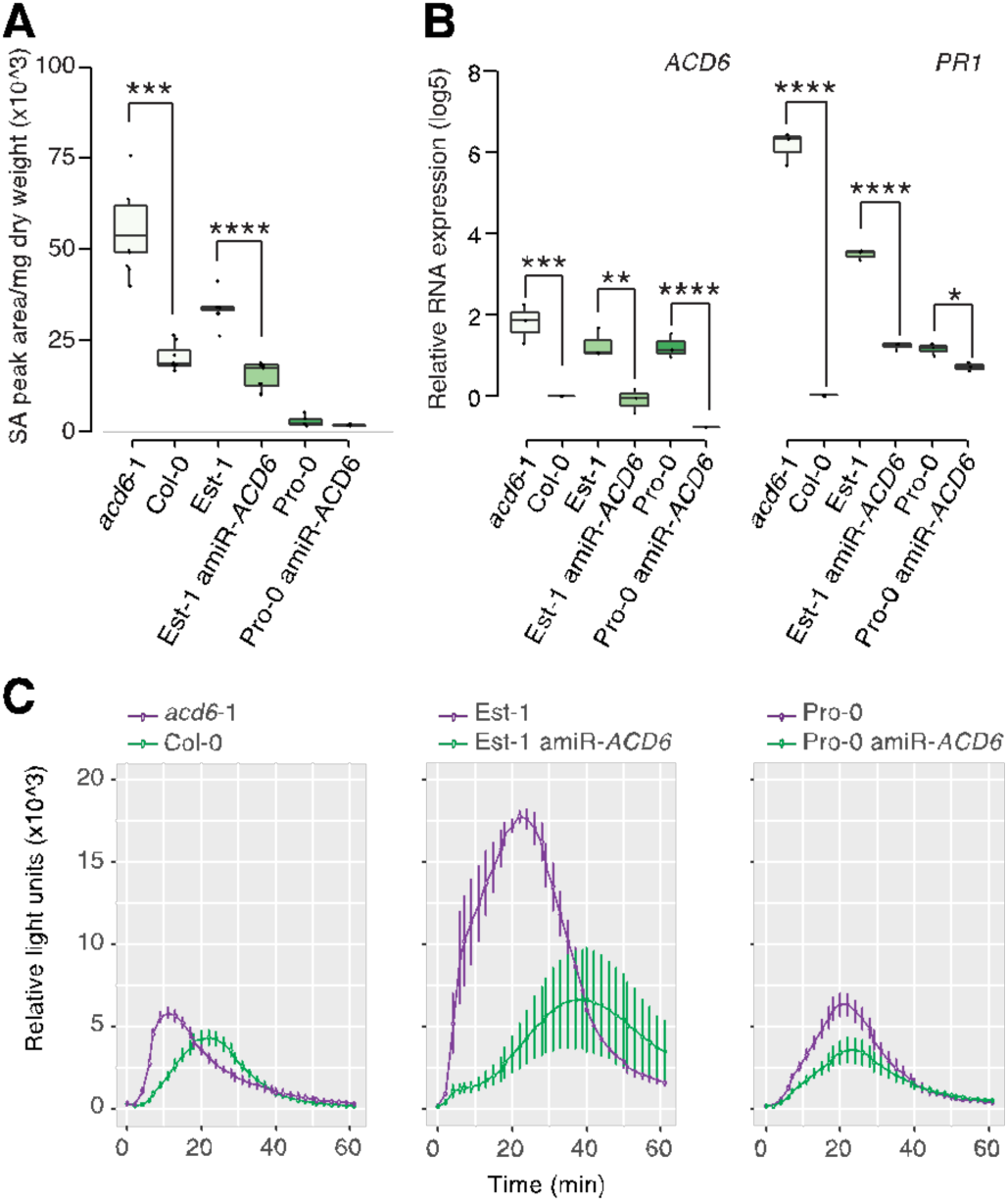
*ACD6*-dependent suppression of salicylic acid accumulation and flg22 responses in Pro-0. (A) SA levels in lines with different *ACD6* alleles, from ten biological replicates each. ^∗∗∗∗^ *p*<0.0001, ^∗^ *p*<0.05 (Student’s t-test). (B) *ACD6* and *PR1* expression in different T_1_ individuals from 4 biological replicates, normalized to Col-0. (C) Reactive oxygen species (ROS) production in response to flg22 (4 biological replicates). Error bars represent standard errors. Experiments were repeated three times with similar results.

### Modulation of *ACD6* hyperactivity by *RPP4/5* alleles

To understand the population genetic architecture underlying extragenic suppression of the activity of *ACD6*-Est-type alleles, we crossed three accessions that carried such alleles, but did not show necrosis, to the necrotic Est-1 accession. The ratios of necrotic and non-necrotic F_2_ individuals suggested - more than one locus in all mapping populations **(S2 Table)**. We used RAD-seq [41] followed by QTL mapping to identify causal regions of the genome in these accessions. As expected, we identified multiple significant QTL in all mapping populations **(Fig 3A).** The QTL explained 4% to 23% of the observed phenotypic variation, further supporting the quantitative nature of phenotypic suppression of *ACD6*-Est alleles **(S3 Table)**.

**Figure 3:**
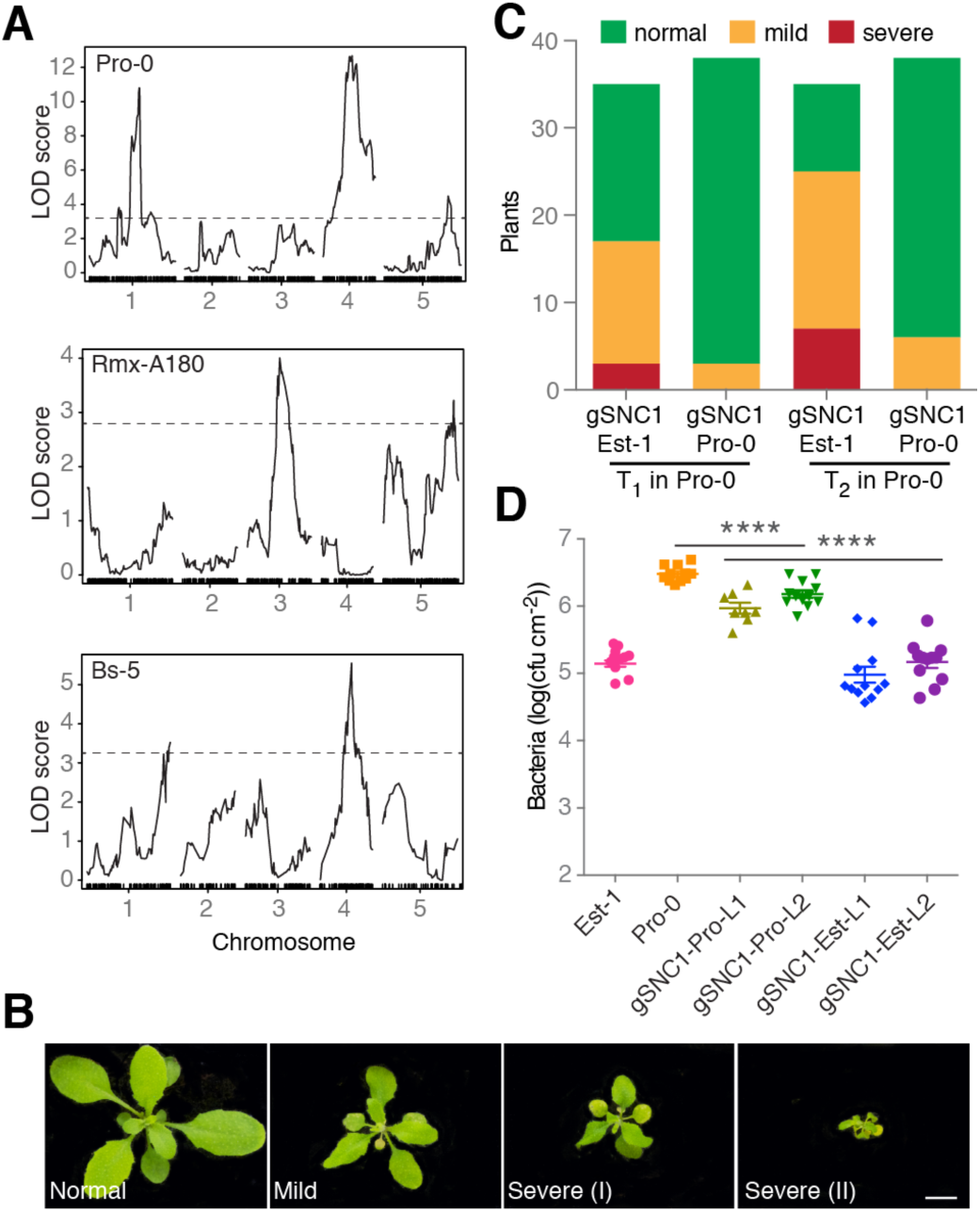
Genetic analysis of *ACD6* modifiers in natural accessions. (A) Necrosis QTL mapping in three F_2_ populations. Dashed lines indicate LOD score significance thresholds of *p*<0.05 (from 1,000 permutations. (B) Representative phenotypes of transformants expressing genomic *SNC1* fragments. Scale bar, 10 mm. (C) Necrosis in transgenic Pro-0 plants. The effects of *SNC1*-Est-1 and *SNC1*-Pro-0 transgenes were significantly different (chi-square test, p<0.001). (D) Growth of Pst DC3000 after infection at OD600 = 0.0001. Bacterial growth in Pro-0 [*gSNC1*-Pro-0] was significantly different from either Pro-0 [*gSNC1*-Est-1] or non-transgenic Pro-0 on day 3. ∗∗∗∗ *p*<0.0001 (one-way ANOVA). L1/L2 designates two independent transgenic lines for both constructs.

We decided to focus on the QTL in the center of chromosome 4, near *ACD6*, which explained 18% of phenotypic variance in the Pro-0 x Est-1 cross, because the confidence interval (Chr 4: 7.5-9.7 Mb, LOD=13.1, p<0.0001) contained a well-studied locus that had been linked to suppression of autoimmunity. Alleles of this locus, *SNC1*, can fully or partially suppress the phenotypic defects of different autoimmune mutants [42,43]. In addition, *SNC1* gain-of-function alleles reduce plant size, similar to *ACD6* hyperactivity, while down-regulation of *SNC1* homologs increases plant size [19,44]. *SNC1* is located in a complex NLR gene cluster, *RECOGNITION OF PERONOSPORA PARASITICA 4/5* (*RPP4/5*), which has alleles that mediate resistance to different strains of the oomycete pathogen *Hyaloperonospora arabidopsidis* [45,46]. *SNC1* and different *RPP4* and *RPP5* alleles are closely related in sequence [19,44,45,47].

To test whether *SNC1* contributes to the suppressed leaf necrosis in Pro-0, and whether *SNC1* can modulate *ACD6* activity, we took a transgene approach. We first introduced a SNC1-Pro-0 genomic fragment into Pro-0. This transgene caused autoimmunity, consistent with additional NLR copies often leading to spontaneous activation of immunity, but the degree of autoimmunity was very modest [48–50]. In contrast, when we introduced a chimeric transgene containing the *SNC1* promoter from Pro-0, but coding sequences from Est-1, into Pro-0, we observed strong necrosis and dwarfism, both in primary transformants and in the T_2_ generation **(Fig 3B and 3C, S1 Fig)**. In agreement, Pro-0 plants transformed with the *SNC1*-Est allele were more resistant to infection by *P. syringae* pv. *tomato* strain DC3000 (*Pto* DC3000) than untransformed Pro-0 plants or Pro-0 plants carrying an extra *SNC1*-Pro copy (*p*<0.0001) (**Fig 3D**). This suggests that differences in the transcribed portion of *SNC1*, most likely differences between the *SNC1*-Est and *SNC1*-Pro proteins, are causal for attenuated effects of *ACD6*-Est alleles.

### Correlation of a common *SNC1* structural variant with attenuation of *ACD6* activity

To gauge whether attenuation of *ACD6*-Est activity by variant *SNC1* alleles is likely to be common, we wanted to assess the population frequency of different *SNC1* alleles. We compared the *SNC1* alleles from Est-1 and Pro-0 with *RPP4/RPP5/SNC1*-like sequences from a REN-seq dataset of 65 *A. thaliana* accessions [51]. A protein-phylogeny of 142 *RPP4/RPP5/SNC1* homologs indicated that 34 genes, from 29 accessions, form a single clade of *SNC1* orthologs, which are predicted to encode highly similar Toll-Interleukin-1 receptor (TIR) domains. Five accessions, Ws-2, Liri-1, Nicas-1, DraIV 6-22 and MNF-Che-2, have two *SNC1* homologs each **(Fig 4A, S2 Fig, S5 Table)**. The most noticeable sequence differences among SNC1 proteins affect the NL linker between the nucleotide binding (NB) and leucine-rich repeat (LRR) domains [19]. The NL linker (amino acids 544-671 in SNC1-Col-0) is duplicated in 15 of the 34 SNC1 proteins **(Figs 4A and 4B, S2 Fig and S3 Fig)**. A *SNC1* allele that differs functionally from the Col-0 reference allele has been identified in Ws-, *SNW* [48]. The *SNC1* homolog that we found in addition to *SNW* in Ws-2 (Ws-2-T078) has a duplicated linker and is highly similar to the Pro-0 variant **(Fig 4B, S4 Fig)**. Similarly, the MNF-Che-2 and DraIV 6-22 accessions have two *SNC1* genes, one each encoding a single and a duplicated linker **(Fig 4B)**. Overall, our analysis revealed a complex history of *SNC1* diversification within and between accessions.

**Figure 4.**
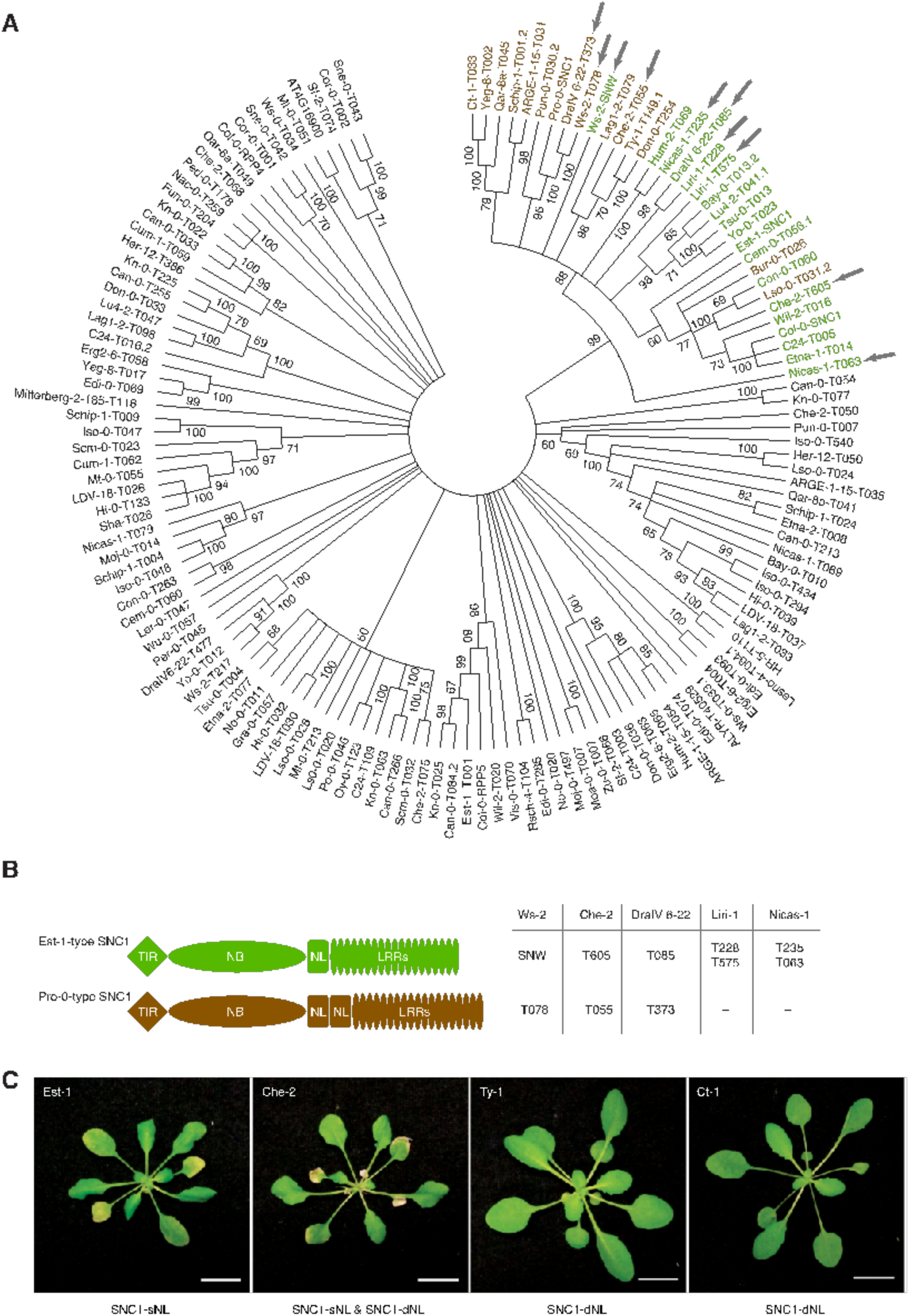
Correlation between *SNC1* alleles and *ACD6*-Est-1 effects in natural accessions. (A) Phylogeny of 142 RPP4/5 and SNC1 proteins from 65 accessions. Bootstrap values over 60% are indicated. The SNC1 clade is highlighted in color, with single NL linker (SNC1-sNL) genes green and duplicated NL linker (SNC1-dNL) genes brown. Arrows highlight five accessions with two *SNC1* homologs. (B) Diagrams of SNC1 proteins with single and duplicated NL linkers. Gene IDs for five accessions with two *SNC1* homologs on the right. (C) Accessions with *ACD6*-Est-like alleles; SNC1 types indicated below. Scale bar, 10 mm.

To address whether SNC1 linker variation is associated with suppression of *ACD6* activity across accessions, we determined the *ACD6* allele type in all 15 accessions with the duplicated SNC1 linker. Three accessions, Ty-1, Ct-1 and MNF-Che-2, had *ACD6*-Est alleles **(S5 Fig)**, but only MNF-Che-2 had symptoms of necrosis. This is in agreement with our hypothesis that weak *SNC1* alleles can function as genetic suppressors of *ACD6* hyperactivity **(Fig 4C)**. That *ACD6*-Est induced necrosis was not suppressed in MNF-Che-2 may be due to the additional *SNC1* homolog without a duplicated linker in this accession **(Fig 4C)**.

### Experimental evidence for SNC1 NL linker affecting *ACD6* activity

A gain-of-function allele of *SNC1*, *snc1*-1, has a single amino acid substitution in the NL linker domain, pointing to the functional importance of the NL linker [19]. Since the NL linkers of SNC1 from Pro-0 and Est-1 differ notably, we suspected that they might be responsible for differential *ACD6*-Est effects in different backgrounds. To test this hypothesis, we introduced three different *SNC1* constructs into Col-0: a genomic *SNC1*-Pro fragment, a genomic *SNC1*-Est fragment, and a chimeric SNC1-Pro fragment with the NL linker replaced with the linker from Est-1. Sixteen of 53 *SNC1-Est* T1 transgenic plants had a severe autoimmune phenotype **(Fig 5A)**. However, none of 70 *SNC1*-*Pro* T_1_ transgenic plants showed severe, and only 10% showed mild phenotypes. The ability of *SNC1-Pro* in triggering leaf necrosis was fully restored when introducing the SNC1-Est linker in an otherwise Pro-0 protein (SNC1-Pro-NL-Est) **(Fig 5A)**. To further assess the contribution of the NL linker on SNC1-induced cell-death signaling, we transiently expressed both SNC1-Est and SNC1-Pro in *N. benthamiana* and monitored chlorosis indicative of cell death 7 days after infiltration. Expression of SNC1-Pro resulted in a much weaker cell death and significantly less ion leakage than either SNC1-Est or SNC1-Pro-NL-Est **(Fig 5B, S6 Fig)**. Taken together, we conclude that polymorphisms in the NL linker region explain functional differences between SNC1-Est and SNC1-Pro.

**Figure 5.**
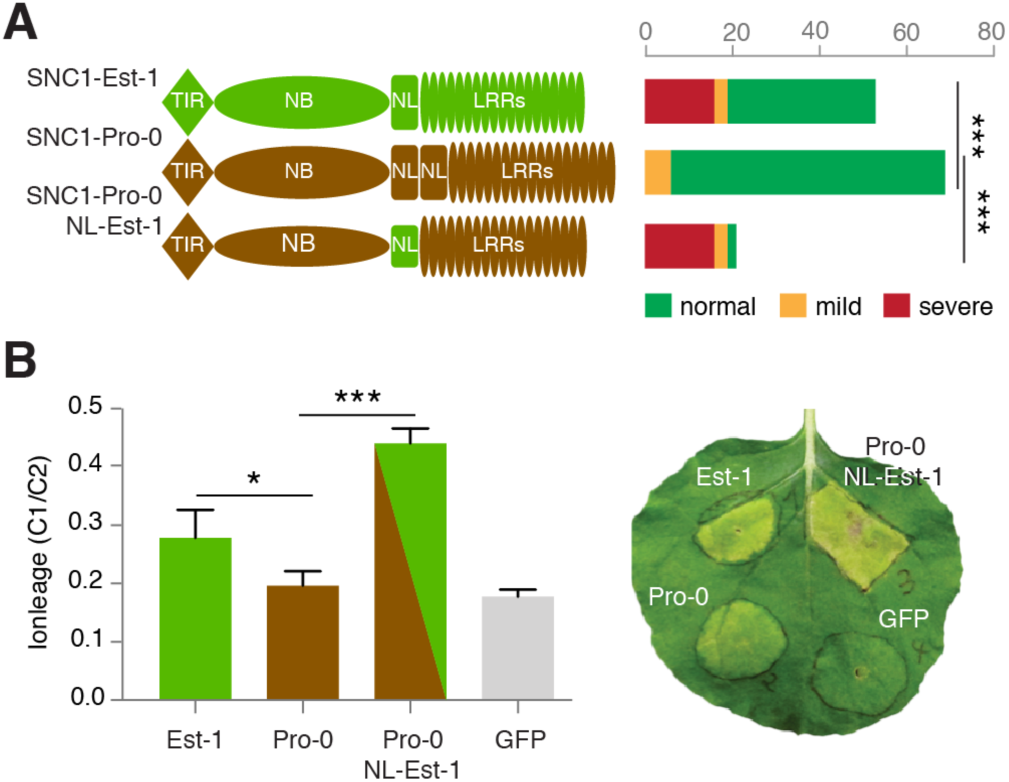
Contribution of NL linker variation to SNC1 activity. (A) Distribution of necrosis in Col-0 T_1_ transformants expressing *gSNC1*-Pro-0 with different NL linkers. ^∗∗∗^, *p*<0.001 (chi-square test). (B) Left, ion leakage, an indicator of cell death, in *N. benthamiana* leaves transiently transformed with different *gSNC1* transgenes 4 d after *Agrobacterium* infiltration. The leaf samples were collected from six independent plants. ^∗^ *p*<0.05, ∗ *p*<0.001 (Student’s t-test). Right, image of a representative leaf 7 d after infiltration.

## Discussion

Darwinian fitness is defined as the number of fertile offspring an individual has [52]. Maximizing fitness requires the right balance among investments in growth, reproduction and defense against biotic and abiotic challenges [2,3,53]. The naturally hyperactive *ACD6*-Est allele in *A. thaliana* shifts the balance from growth to defense in many accessions, but this does not appear to be the case for all accessions with this allele [32]. We have shown that variation in the effects of *ACD6*-Est alleles does not map to the *ACD6* locus itself, but rather is caused by extragenic suppressors of the *ACD6*-Est phenotype. Notably, alleles at several different loci can have such an effect, including a common allele at the NLR locus *SNC1.*

Genetic mapping experiments with three different accessions carrying an *ACD6*-Est allele identified a minimum of six genomic regions that can attenuate *ACD6*-Est effects. The identified QTL only explain a minority of observed phenotypic variation, suggesting that several additional minor-effect loci contribute to modulation of *ACD6* activity. An attractive hypothesis is that suppressors of *ACD6*-Est-like activity have arisen as a consequence of excessive fitness costs of *ACD6*-Est and perhaps hyperimmunity alleles at other loci. Alternatively, the *ACD6*-Est allele itself may first have evolved on a background that masked its effects. We found that about a quarter of *A. thaliana* accessions share the *SNC1* allele with a duplicated NL linker, which can attenuate *ACD6*-Est effects. The fraction of accessions with an *ACD6*-Est allele among those with an *SNC1* allele is similar to the frequency of *ACD6*-Est alleles in the global *A. thaliana* population. While there is no obvious indication for linkage between specific *SNC1* and *ACD6* alleles, the numbers examined so far are very small, and the situation could be different in local *A. thaliana* populations. In addition, any claims about co-evolution of *ACD6* and other loci must take other natural modifiers of *ACD6*-Est alleles into account.

It has been argued that tandem duplication of NLR genes within clusters can provide an effective mechanism to acquire novel NLR specificities without sacrificing old ones [54]. *SNC1* features both relevant structural and copy number variation, and functionally distinct variants can occur in the same genome, offering an opportunity to test such hypotheses about NLR evolution. Since *SNC1* has a dosage effect on plant innate immunity [48], it will be interesting to ask what fitness costs are associated with having two functionally different *SNC1* copies, as we have seen in MNF-Che-2, Ws-2 and DraIV 6-22.

In summary, our genetic analyses of accessions with a hyperactive *ACD6* allele have shown that natural *A. thaliana* populations frequently harbor extragenic modulators of *ACD6* activity, underscoring the importance of epistatic interactions for natural phenotypes. In addition, our work has revealed a new connection between *ACD6*, which has before been primarily linked to PAMP recognition and PTI [30,31,55], and NLR proteins, which are central to ETI. There is already genetic evidence for cell death triggered by PAMP receptor complexes being suppressed by the NLR SNC1 [48,56]. An arbitrary selection of evidence for the distinction between PTI and ETI being fluid [35] includes the overlap in the signaling network downstream of AvrRpt2-triggered ETI and the network activated by the well-known PAMP flg22 [57], or MAPK3 and MAPK6 as widely shared downstream signaling components for both PTI and ETI [57–59]. Together, our findings illustrate the potential of studies of naturally occurring autoimmunity in either *A. thaliana* inbred strains or hybrids [60] to contribute to our understanding of the immune system in healthy and diseased plants.

## Methods

### Plant material and growth conditions

Seeds of *Arabidopsis thaliana* accessions **(S1 Table and S5 Table)** were from stocks available in the lab. Seeds were germinated and cultivated in growth rooms at a constant temperature of 23°C (temperature variability about ± 0.l°C), air humidity of 65% and long days (LD, 16 hr day length) or short days (SD, 8 hr day length), with light (125 to 175 □mol m^-2^ s^-1^) provided by a 1:1 mixture of Cool White and Gro-Lux Wide Spectrum fluorescent lights (Luxline plus F36W/840, Sylvania, Erlangen, Germany). Leaf necrosis was scored in SD at 23°C.

### qRT-PCR

RNA was extracted from three biological replicates (4- to 5-week-old entire rosettes) using TRIzol Reagent (Thermo Scientific, Waltham, MA) and treated with DNase I (Thermo Scientific, Waltham, MA, USA). 2 μg total RNA was used as a template for reverse transcription (M-MLV reverse transcriptase kit; Thermo Scientific, Waltham, MA, USA). Quantitative real-time PCR reactions were performed using Maxima SYBR Green Master Mix (Thermo Scientific, Waltham, MA, USA) according to the manufacturer’s directions on a CFX384 real time instrument (Bio-Rad, Hercules, CA). Transcript abundance was normalized to the *TUBULIN BETA CHAIN 2* (At5g62690) transcript. Relative expression compared to the control (usually Col-0) was quantified based on [61]. Primers used for qRT-PCR are listed in **S7 Table**.

### ROS assay

Leaf discs (5 mm diameter) were punched from 5-week-old 23°C SD-grown plants and immediately floated with the adaxial side up in individual wells of a 96-well plate (Greiner Bio-One, Frickenhausen, Germany) containing 200 □L of water. They were incubated overnight (12-16 hr) covered with a transparent lid. The water was replaced with the elicitation solution containing 10 μM Luminol (Sigma-Aldrich, MO), 10 μg mL^-1^ horseradish peroxidase and 100 nM flg22 (QRLSTGSRINSAKDDAAGLQIA, >85% purity; EZBiolab, Westfield, IN [62]), and ROS production was monitored by luminescence beginning immediately for at least 90 minutes (Tecan Infinite PRO multimode reader, Tecan, Männedorf, Switzerland). Controls were mock treated with the same solution without flg22. Each genotype was assayed at least three independent times.

### Salicylic acid (SA) quantification

Each sample consisted of 10 (± 1) mg freeze-dried, ground plant material of 4-week-old rosettes, with 8 biological replicates per genotype, of plants that had been grown in randomized complete block design at 23°C SD. SA was extracted twice with 400 μl 20% methanol (LCMS-grade) / 0.1% hydrofluoroalkane by 5 min ultrasonic extraction, followed by 20 min incubation on ice, and removal of solids by centrifugation for 10 min at 13,500 g. Ultra Performance Liquid Chromatography Mass Spectrometry (UPLC-MS) analysis was performed on a Waters Acquity UPLC system coupled to a SYNAPT G2 QTOF mass spectrometer equipped with a Zspray TM ESI-Source (Waters Corporation, Milford, MA) at the University of Tübingen ZMBP Analytics Laboratory. MassLynx v4.1 and TargetLynx (Waters Corporation) were used to control the LCMS system and to perform data integration.

### RAD-seq and whole genome genotyping

RAD-seq library preparation using a set of 192 adaptor sequences containing PstI and Mse1 recognition sequences were prepared according to [41,63]. Ninety six or 192 individual samples were pooled per library and sequenced on one Illumina HiSeq 2000 lane (100 bp single-end reads). Raw reads were processed with the SHORE (https://sourceforge.net/projects/shore) pipeline. Default parameters were used for de-multiplexing, read trimming and mapping. The BWA [64] option, also with default parameters, was used to map the reads to the TAIR9 Col-0 reference genome. In SHORE, the consensus sequences were computed, which were used to create a matrix containing the genotypes at specific marker positions for all F_2_ individuals **(S4 Table).** We only considered bi-allelic SNPs that agreed with SNPs from both parents and the F_1_ hybrid of a given cross. Scripts are available at https://github.com/mzaidem/ACD6_suppressors_mapping_RADseq_scripts/tree/master.

### QTL mapping and analysis

Pro-0 x Est-1, Rmx-Al80 x Est-1 and Bs-5 x Est-1 F_2_ individuals were phenotyped for late-onset necrosis in a binary manner. QTL mapping and testing for QTL effects and interactions were performed using the R/qtl package [65]. Lod scores were calculated under a single-QTL model using the function “scanone”. When more than one locus was detected, re-computation of lod scores and interaction testing (epistasis, additive and dominant) was performed using the functions “scantwo” and “fitqtl”. Lod score significance thresholds were established using 1,000 permutations. Bayesian credible intervals were estimated for individual QTL. The standard expectation-maximization algorithm was always used as “method.”

### Transgenic lines

*ACD6* fragments were amplified from genomic DNA with PCR primers designed based on *ACD6* from Est-1 ([32], **S7 Table**). The Pro-0 (Rmx-A160) genomic DNA fragment was 9.1 (10.3) kb long, including 2.5 (1.8) kb sequences upstream of the initiation codon and 2.8 (3.2) kb downstream of the stop codon. Restriction enzyme sites of Gibson Assembly (NEB, Ipswich, the USA) were used to generate chimeric constructs [66]. An amiRNA against *ACD6* (TTAATGGTGACTAAAGGCCGT) [32] was used to knock down *ACD6.* All constructs were individually cloned into the Gateway entry vector pCR8/GW/TOPO (Invitrogen) and moved into binary vector pFK206 by LR reaction (Thermo Scientific, Waltham, MA, USA). Constructs in pFK206 were transferred to *Agrobacterium tumefaciens* strain ASE and plants were transformed by floral dipping [67] **(S6 Table)**.

### Phylogenetic analysis of RPP4/RPP5/SNC1 protein sequences

SNC1 (AT4G16890), RPP4 (AT4G16860), and RPP5 (AT4G16950) like protein sequences from 63 accessions were extracted from an in-house NLR database of Araport II sequences and the single RPP4/5 homolog of *Arabidopsis lyrata* were used as tempeplate to query the database with exonerate v2.2.0 allowing a minimal mapping score of 95% [69]. . We used MEGA6 [70] to realign these sequences with SNC1-Est-1, SNC1-Pro-0, SNC1-Ws-2 (SNW, GenBank accession AY510018), RPP5-Ler-0 (AF180942) and the RPP4/RPP5/SNC1 homolog At4g16900 from Col-0 (Araport 11). One-hundred forty-two RPP4/5 protein sequences were used to infer phylogenetic relationships with both Neighbor-Joining (NJ) and Maximum likelihood (ML) approaches in MEGA6, either over their entire lengths or only the TIR domain (positions 1-226 in SNC1-Col-0). Node confidence was estimated with 1,000 bootstrap replications. All sequences that we could confidently assign to the SNC1 clade were aligned separately to identify functional polymorphisms. A similar approach was used to generate a phylogenetic tree of ACD6 protein sequences extracted from the in-house NLR database, using ACD6 (AT4G 14400) and the adjacent ACD6 homolog AT4G 14390 from Araport11 as references.

### Bacterial infection

*Pseudomonas syringae* pv tomato strain DC3000 was grown to OD_600_ around 3.0, collected and resuspended in 10 mM MgCl_2_ at 5 × 10^5^ colony-forming units (cfu)/ml. The bacterial suspension was infiltrated into 4-week-old leaves with a needle-less syringe. Bacterial growth was determined 3 days post inoculation (dpi).

### Transient expression in *N. benthamiana* and ion-leakage measurements

*Agrobacterium tumefaciens* with binary plasmids was grown to OD_600_ of 1.6, and incubated in induction medium (10 mM MES pH 5.6, 10 mM MgCl_2_ and 150 □M acetosyringone) for three hours.

Cell suspensions were normalized to OD_600_ of 0.8 for co-infiltration into the abaxial side of leaves of *N. benthamiana* leaves that had been grown under SD at 23°C. Images were taken 7 days after infiltration. Conductivity assays to measure ion leakage were performed according to [47]. Six independent leaves were infiltrated with *A. tumefaciens* with binary constructs encompassing different *SNC1* alleles and two discs (0.6 mm diameter) were harvested per leaf 5 days after infiltration. The fresh leaves were briefly rinsed with ddH_2_O and placed in a tube with 8 ml ddH_2_O. Samples were gently shaken for 24 hr before measuring ion content (C1) using an Orion Conductivity (Thermo Scientific, Waltham, MA, USA) instrument. Total ion content (C2) was determined from leaf samples that had been boiled for 15 min. C1/C2 ratios were calculated as indicators of ion leakage. *Nicotiana benthamiana* leaves expressing GFP only were used as control. Statistic differences were calculated by one-way-ANOVA.

## Acknowledgments

We thank the members of the GBMF-funded Ren-seq Team, Freddy Monteiro, Jeff Dangl, Oliver Furzer, Kamil Witek and Jonathan Jones, for their help in generating the Ren-seq NLR database. We thank Chang Liu and the Weigel lab for comments.

## Supplementary Information

**S1 Figure.**
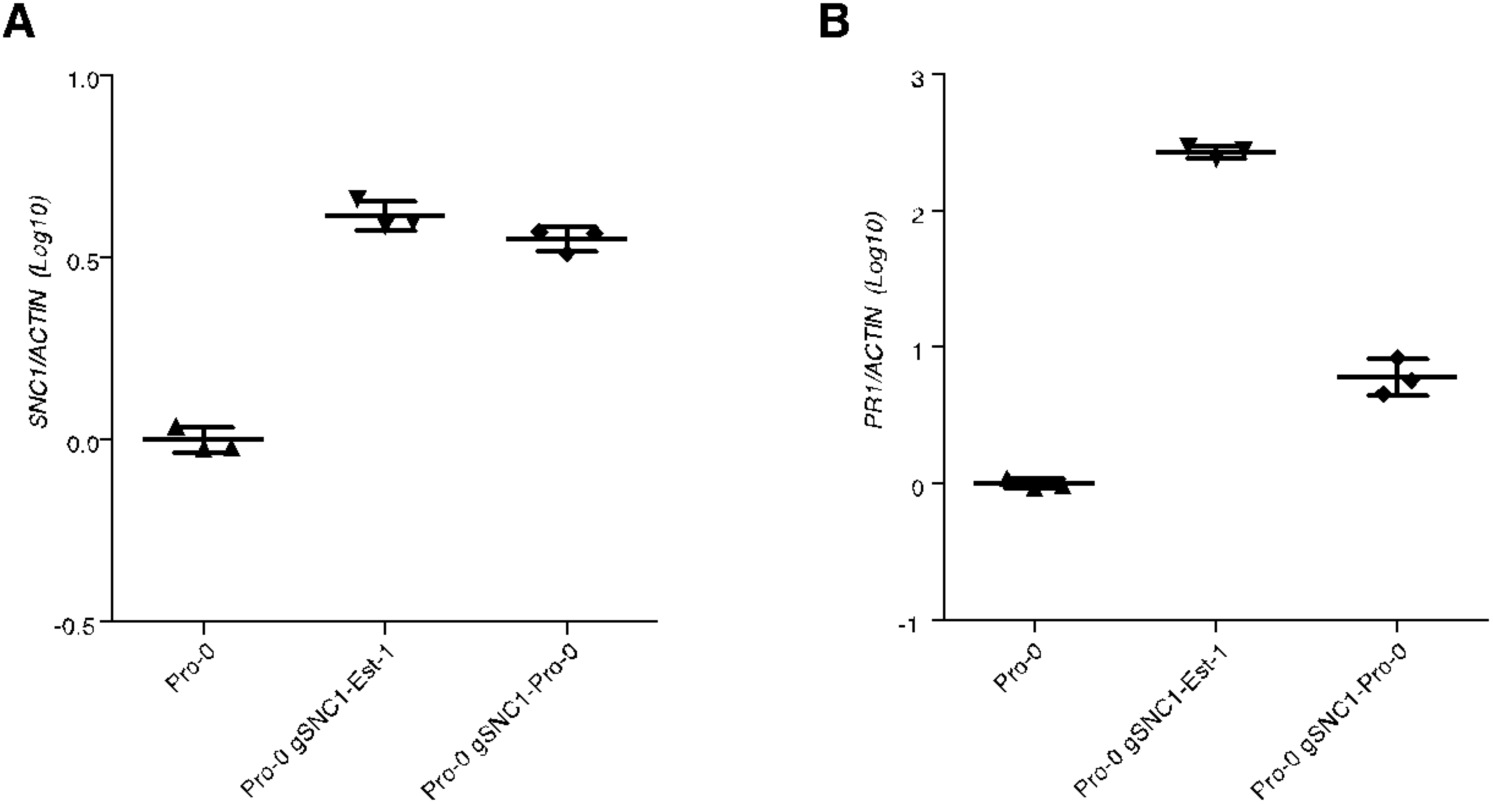
*SNC1* and *PR1* expression in Pro-0 T_2_ transformants, as measured by qRT-PCR. *gSNC1*-Pro-0 transformants with mild necrosis were analyzed.

**S2 Figure.**
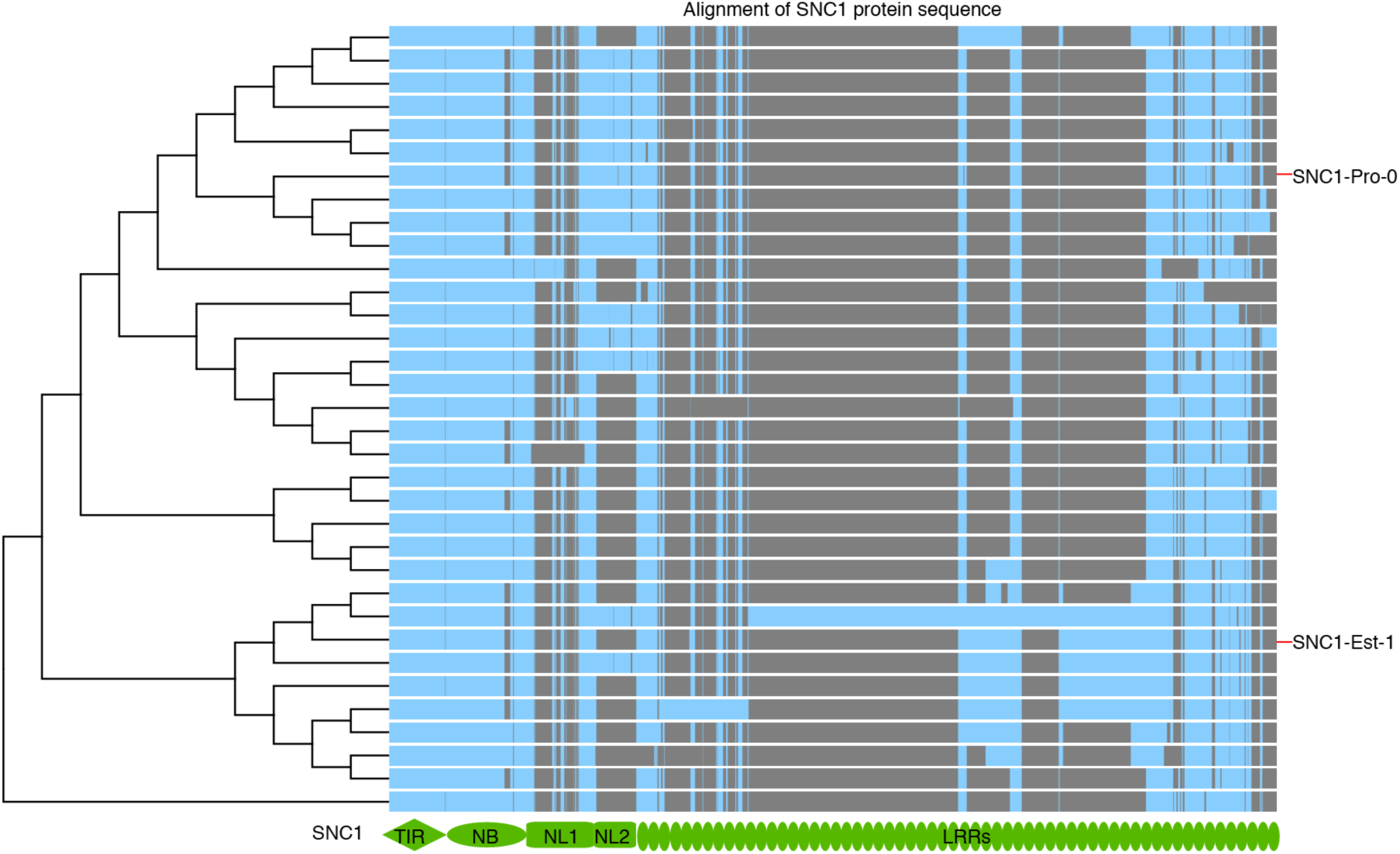
Alignment of 34 SNC1 protein sequences from 29 accessions. Gray indicates regions with structure variation including small and large deletions.

**S3 Figure.**
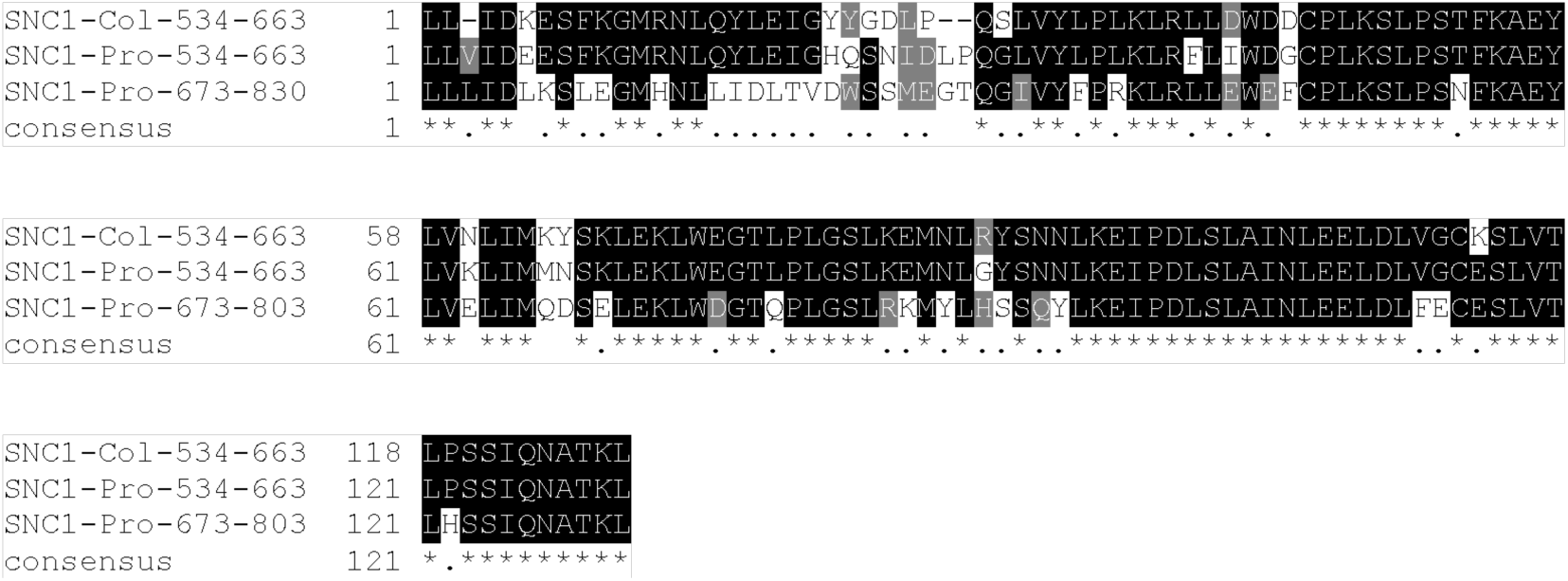
Alignment of duplicated NL linker from SNC1-Pro-0, with SNC1-Col-0 linker as reference.

**S4 Figure.**
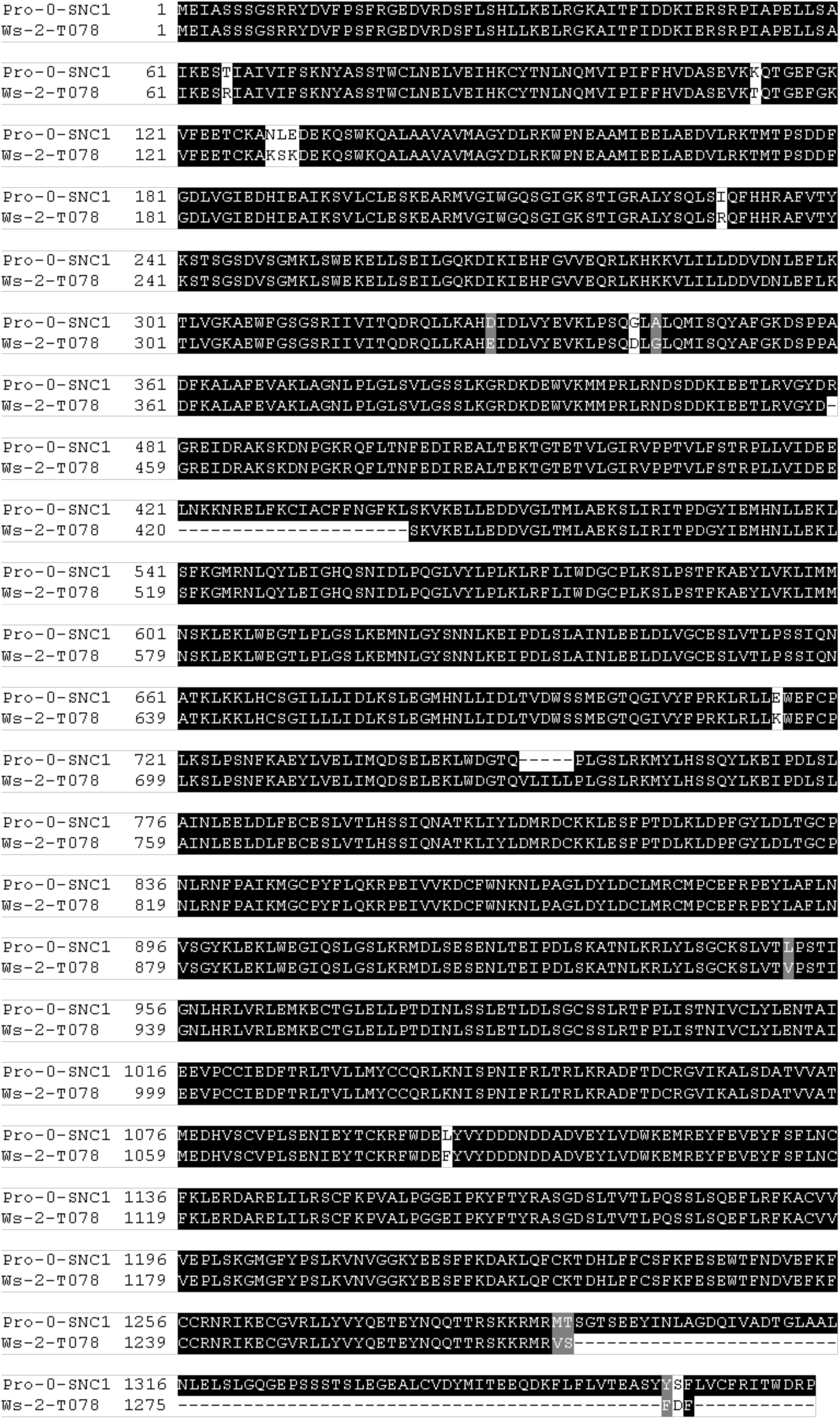
Alignment of Pro-0 and Ws-2 SNC1 homolog. Ws-2-T078 is a newly identified SNC1, in addition to the previously identified loss-of-function SNW variant in Ws-2.

**S5 Figure.**
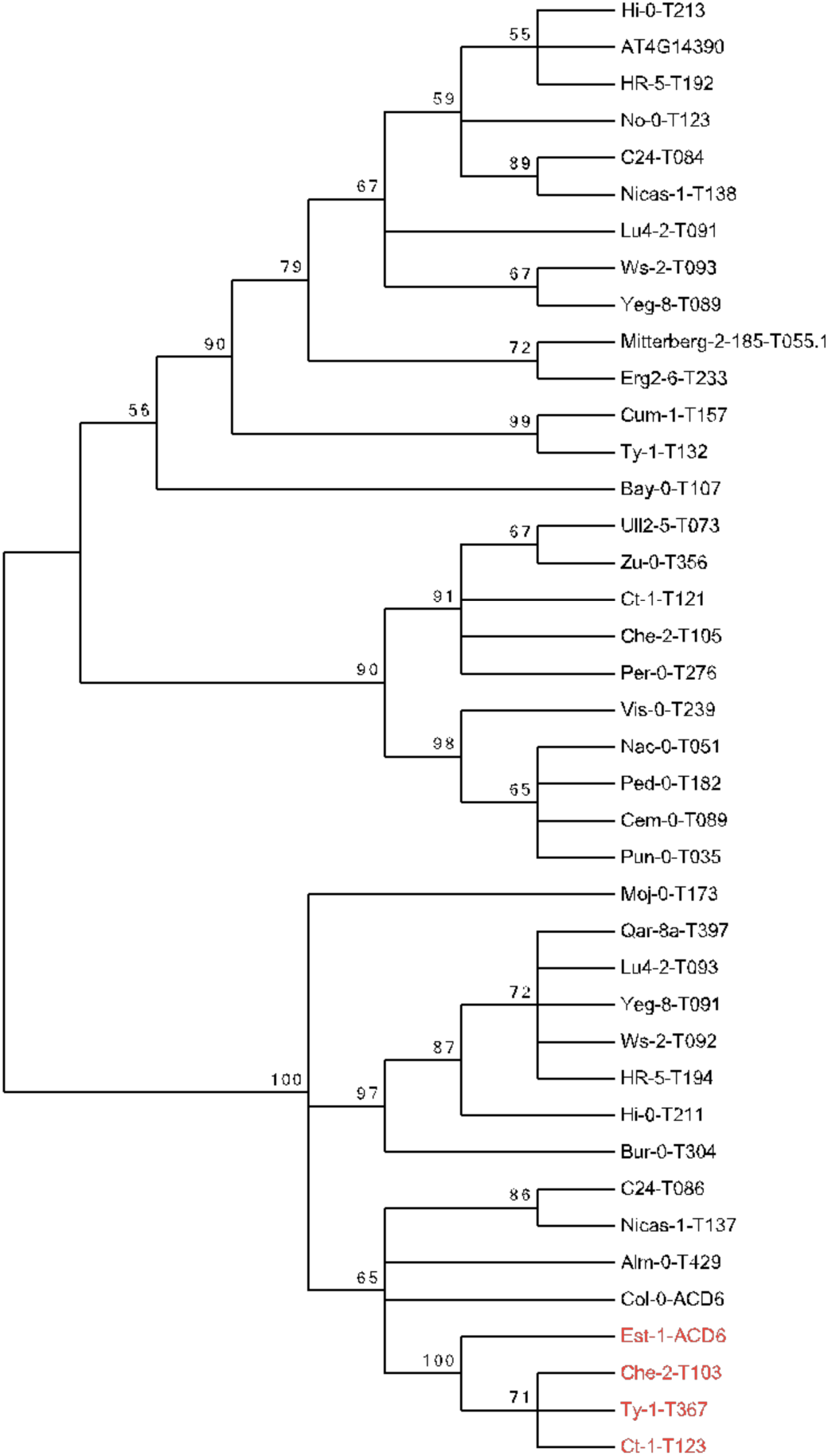
Phylogeny of ACD6 proteins identified from RenSeq. Bootstrap values over 60% are indicated. Est-1-like clade highlighted in red.

**S6 Figure.**
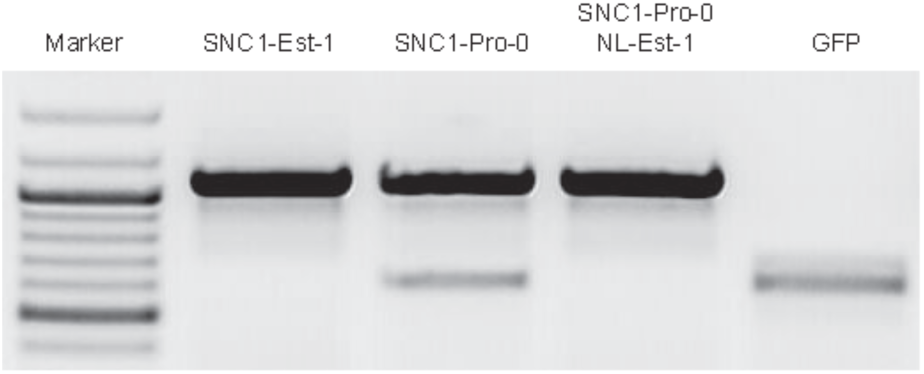
Expression of *SNC1* in infiltrated *N. benthamiana* leaves, as measured by RT-PCR. Samples were harvested 2 d after infiltration.

**S1 Table.** Variation of necrotic phenotype from natural populations with Est-1 like *ACD6* allele.

**S2 Table.** Summary of phenotypic variance explained given a full, additive and interaction model between QTL 1 and QTL 2 of each mapping population used for QTL mapping.

**S3 Table.** Summary of phenotypic variance explained taking individual QTLs separately for each mapping population used for QTL mapping.

**S4 Table.** Genotypes of three F_2_ mapping populations derived from Est-1 and Pro-0, Rmx-A180, and Bs-5.

**S5 Table.** Accessions in phylogenetic tree of RPP4/RPP5/SNC1.

**S6 Table.** Binary T-DNA constructs.

**S7 Table.** Primers used for generating constructs and qRT-PCR.

